# Interactions between culturable bacteria are highly non-additive

**DOI:** 10.1101/2022.09.01.506178

**Authors:** Amichai Baichman-Kass, Tingting Song, Jonathan Friedman

## Abstract

Microorganisms are found in diverse communities whose structure and function are determined by interspecific interactions. Just as single species seldom exist in isolation, communities as a whole are also constantly challenged and affected by external species. Though much work has been done on characterizing how individual species affect each other through pairwise interactions, the joint effects of multiple species on a single (focal) species, remain under explored. As such, it is still unclear how single species effects combine to a community-level effect on a species of interest. To explore this relationship, we assayed over 14,000 communities of two, three, and four bacterial species, measuring the effect of single, pairs of, and trios of 61 affecting species on six different focal species. Our results demonstrate that joint effects of multiple species on a focal species are typically not given by the sum of the effects of individual affecting species. Rather, they are dominated by the strongest individual-species effect. Therefore, while joint effects of multiple species are often highly non-additive, they can still be derived from the effects of individual species, making it plausible to map complex interaction networks based on pairwise measurements. This finding is important for understanding the fate of species introduced into an occupied environment, and is relevant for applications in medicine and agriculture, such as probiotics and biocontrol agents, as well as for ecological questions surrounding migrating and invasive species.

## Introduction

Scarce are the environments on Earth not colonized by bacteria. In addition to naturally playing important roles from driving biogeochemical cycles at the ecosystem level (Cavicchioli et al. 2019; Arrigo 2005; Falkowski, Fenchel, and Delong 2008) to supporting host health at the individual level (Berendsen, Pieterse, and Bakker 2012; Manor et al. 2020; Gilbert et al. 2018), bacteria have also been harnessed for countless biotechnological applications across industries such as food preservation (Motarjemi 2002), plant and animal health (Berendsen, Pieterse, and Bakker 2012; de Mello Júnior et al. 2021), biocontrol of pathogens (Köhl, Kolnaar, and Ravensberg 2019), as well as decomposition of toxic compounds, and production of valuable ones (Varjani, Gnansounou, and Pandey 2017; Ro et al. 2006; Fang and Smith, 2016; Mainka et al. 2021). In natural environments bacteria often form rich and complex communities, but understanding how these communities organize has proven difficult (Widder et al. 2016). Elucidating the rules that govern microbial ecology can both offer insight into larger ecological systems, and allow us to better manipulate and design microbial communities for desired functions.

The structure of microbial communities is determined by the interactions between the involved species (Konopka, Lindemann, and Fredrickson 2015; Barbier et al. 2018; Qian and Akçay 2020). In recent years, much effort has been put into measuring pairwise interactions of different species from, and in, different environments (Foster and Bell 2012; Vetsigian, Jajoo, and Kishony 2011; Kehe et al. 2021). But it is still unclear to what extent the joint effects of multiple species on a focal species of interest (e.g. a pathogen) can be inferred from pairwise measurements. Such inference may be challenging due to the presence of indirect interactions: the affecting species may alter each other’s abundances (termed interaction chains), or may modify each other’s effect on the focal species (termed interaction modification, or higher-order interactions) (Sanchez 2019; Wootton 2002).

Despite a strong theoretical foundation, empirical studies in recent years have shown conflicting results regarding the importance of higher order interactions and indirect effects (Levine et al. 2017). For some functions, such as degradation of complex molecules (Sanchez-Gorostiaga et al. 2019; Gralka et al. 2020) and antibiotic production (Tyc et al. 2014; Qi et al. 2021; Westhoff et al. 2021), clear evidence of such interactions has been found. Furthermore, in both empirical and theoretical studies, the presence or absence of an additional species has been shown to affect interactions, and even the outcome of invasion and coexistence in some systems (Mickalide and Kuehn 2019; Chang et al. 2022; Hsu et al. 2019). Additional theoretical work has shown that commonly used ecological models (i.e. generalized Lotka-Voltera), don’t properly capture microbial community interactions, partially due to the nature of these interactions (chemically mediated as opposed to predator-prey) (Momeni, Xie, and Shou 2017). However other studies have shown that both structures of, and interactions within, larger communities can be accurately predicted from pairwise interactions alone, using variations of said models (Friedman, Higgins, and Gore 2017; Meroz et al. 2021; Guo and Boedicker 2016; Os et al. 2018). This being the case, how microbial interactions combine into the joint effect of multiple species on a single species of interest is still poorly understood.

In our research, we used high-throughput nanodroplet-based microfluidics to measure over 14,000 bacterial communities composed of subsets of a library of 61 soil and leaf isolates of which six were fluorescently labeled (Fig. S1). We quantified the effect of individual species and the joint effects of species pairs and trios on the growth of six focal bacterial species and found that the effect of multiple species combine non-additively and are heavily dominated by the strongest single-species effect.

## Results

We conducted high-throughput assays involving 61 affecting species and six different focal species to understand the effects of single species, pairs, and trios on the growth of a given (focal) species. The 61 affecting species included soil and leaf isolates as well as lab strains representing 19 genera from four phyla: Proteobacteria (n=14), Firmicutes (n=2), Bacteroidetes (n=2), and Actinobacteria (n=1) (full list in SI Table 1 and Data S1). The focal species were a subset of six of these species (all proteobacteria) that were transformed to constitutively express a fluorescent protein: (*Escherichia coli* [EC], *Ewingella americana* [EA], *Raoultella planticola* [RP], *Buttiauxella izardii* [BI], *Citrobacter freundii* [CF], and *Pantoea agglomerans* [PA]). Except for EC which is a lab strain (*E. coli K-12* substr. *MG1655*), all focals were isolated from soil samples (Kehe et al. 2021). First, we characterized each species phylogenetically by performing Sanger sequencing of their 16S ribosomal RNA gene, and phenotypically by growing each species on each of 20 different carbon sources and 11 antibiotics. The species showed large variability in carbon utilization profiles with no species growing well on all carbon sources (Fig. S2). There was also high variability in growth on antibiotics with 15 species showing little or no growth on any antibiotics, while 16 species showed resistance to at least 7 antibiotics (Fig. S3).

We performed the interaction assays in the kChip microfluidics device (Kulesa et al. 2018; Kehe et al. 2019), allowing for extensive screening in parallel (see Materials and Methods, Fig. S1). We measured the effects of 354 single species (either 58 or 59 for each of the 6 focal species), the joint effects of 9988 species pairs (between 1571 and 1762 for each of the 6 focal species), and the joint effects of 3,755 species trios (from a subset of 30 affecting species on one focal species). Interaction assays were carried out in minimal M9 media with 0.5% [w/v] glucose for 24 hours. The growth of the focal species was measured by fluorescence, and effects were calculated as the log ratio of growth in coculture to growth in monoculture (Fig. 1D,). Positive and negative effects are defined as a net increase or decrease in growth compared to the monoculture respectively, while affecting species with no observable effect (see Materials and Methods) were defined as neutral.

**Figure 1:**
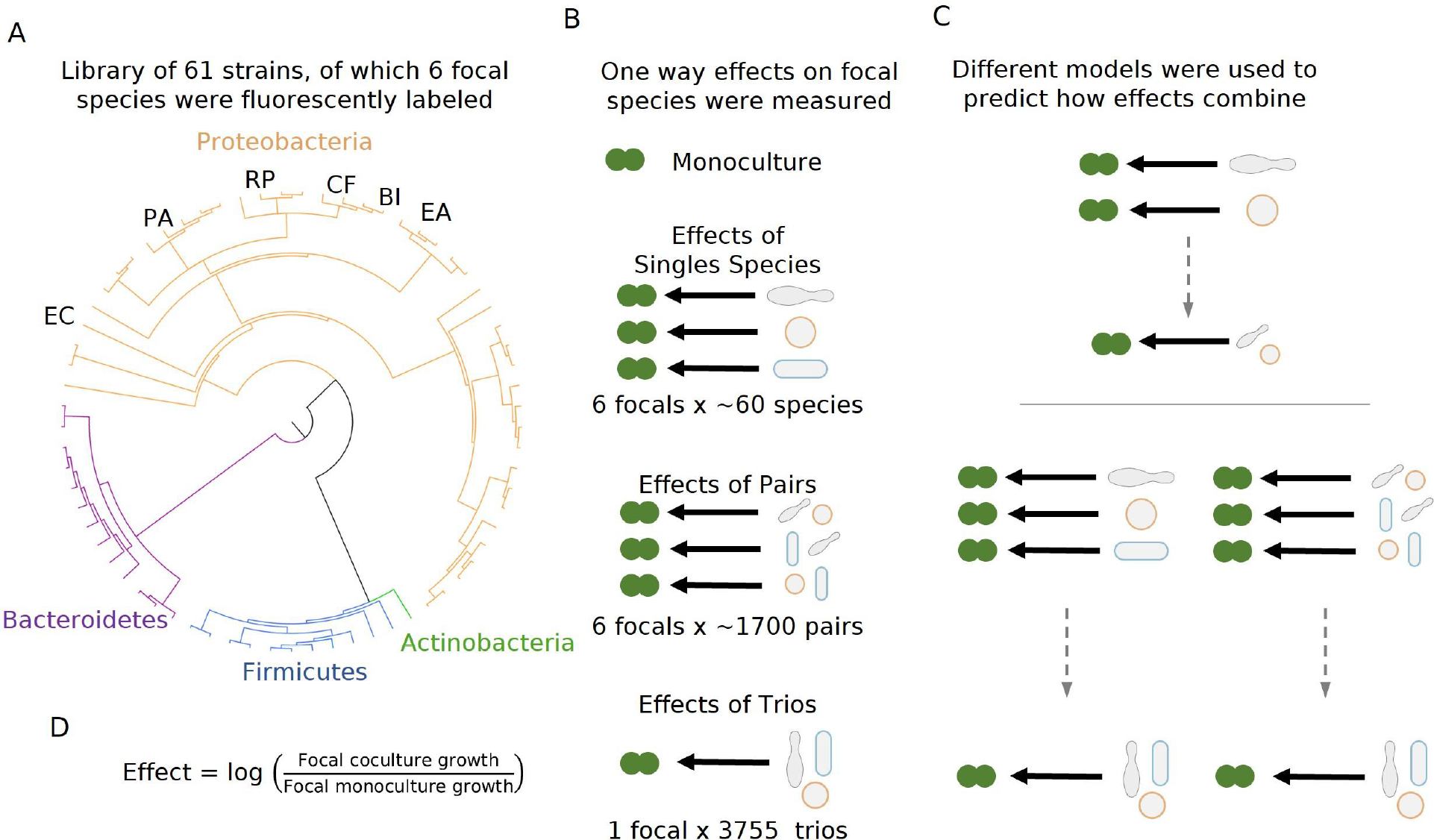
Measuring effects of 61 affecting species, and their pairs and trios on 6 focal species. A) A library of 61 soil and leaf-associated bacterial strains was used in this experiment. All strains are from 4 orders: Proteobacteria (orange), Firmicutes (blue), Bacteroidetes (purple), and Actinobacteria (green) (full list in table S1, Data S1). Six of the 61 species were labeled with GFP and used as ‘focal’ species whose growth was tested in the presence of the other isolates (affecting species). These strains are labeled on the on the phylogenetic tree (Escherichia coli [EC], Ewingella americana [EA], Raoultella planticola [RP], Buttiauxella izardii [BI], Citrobacter freundii [CF], and Pantoea agglomerans [PA], B) Each focal species was grown in monoculture, with (between 58-59) single affecting species, and (between 1571-1762) pairs of affecting species. Additionally, E.coli was grown with 3,755 trios of affecting species. C) Effects of pairs and trios were then predicted using the effects of single species and single species and pairs respectively. Predictions were made using three different models: Additive, Mean, and Strongest (detailed in Results and Materials and Methods). D) Equation used for calculating the effect of an affecting species on the focal species.

### Joint effects of species pairs tend to be more negative than those of individual affecting species

We started our interaction assays by measuring the individual effects of single affecting species on each of the focal species (see Materials and Methods). Individual effects covered a wide range (median=−0.17, interquartile range=0.98) (Fig. 2A), and positive effects (the focal species reaching a higher density in the presence of an affecting species than in monoculture) were common overall (34.3%, Fig. 2B), in line with previous studies(Kehe et al. 2021). The distribution of effects varied based on the focal species, with *E. coli* and *B.izardii* showing the most negative (−2.27) and positive (+0.19) median effects, respectively (Fig. 2D). Additionally, we found no affecting species that had strong effects across all focal species (Fig. S4).

**Figure 2:**
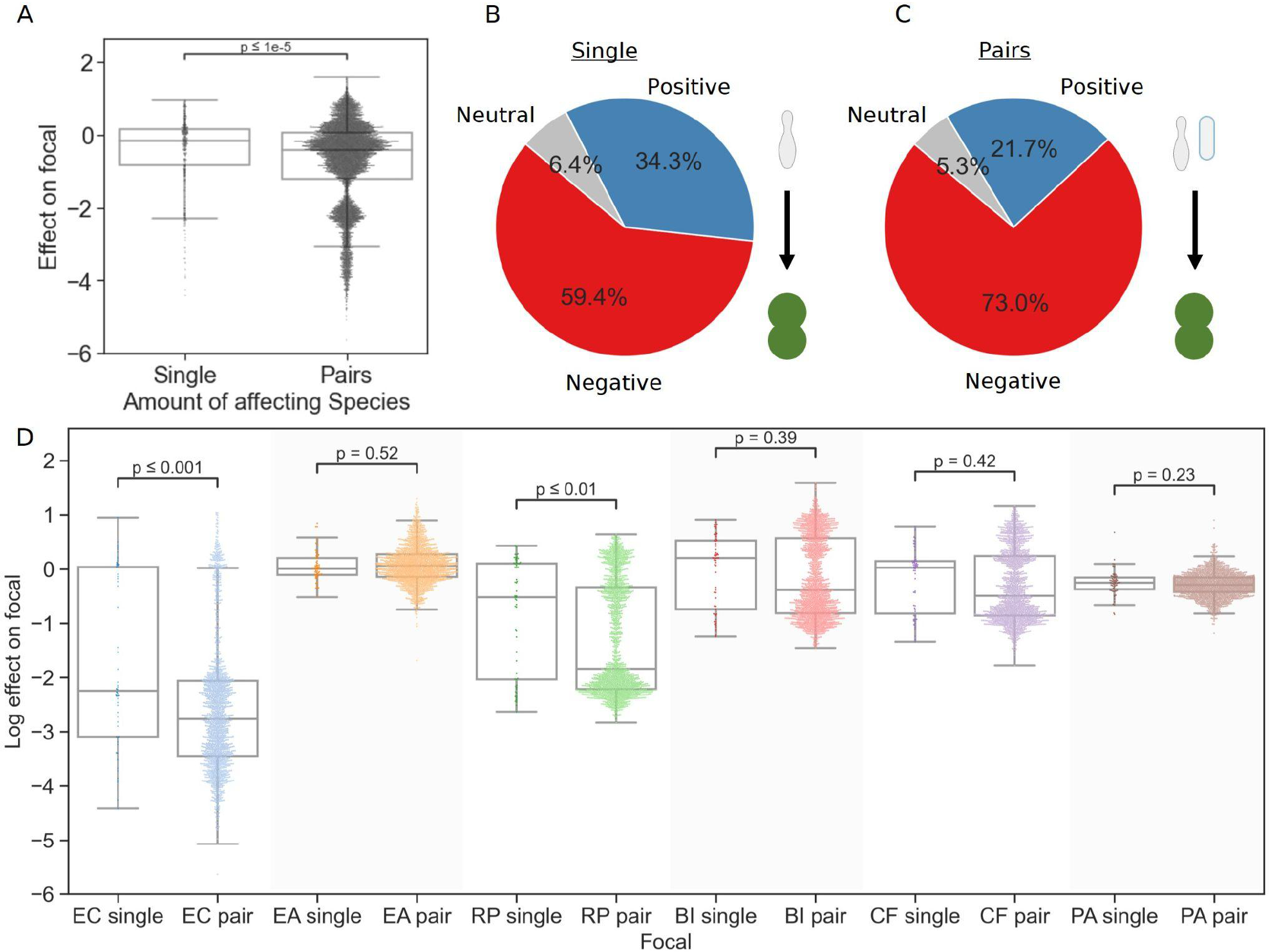
Pairs of affecting species have more negative effects than single species. A) Distribution of the effects of single and pairs of affecting species on all focal species. Mann-Whitney-Wilcoxon test two-sided, p-value=1.6e-10. Dots show individual effects, solid lines represent the median, boxes represent the interquartile range, and whiskers are expanded to include values no further than 1.5X interquartile range. B,C) Distribution of qualitative effects of single and pairs of affecting species respectively on all focal species. D) Distribution of the effect of single and pairs of affecting species for each focal species individually. Dots represent individual measurements, solid lines represent the median, boxes represent the interquartile range, and whiskers are expanded to include values no further than 1.5X interquartile range. Mann-Whitney-Wilcoxon test two-sided tests were performed for each focal species, and p-values are shown on the graph.

The measured traits of individual species showed no consistent correlations with their effects on the focal species. In particular, the similarity of metabolic profile, resistance profile, or phylogeny between the focal and affecting species did not correlate strongly with the effect across focal species. Some traits showed little to no correlation for most focals (e.g. antibiotic resistance), while other traits were correlated with effect for a number focal species but not all (e.g. phylogenetic distance). Most of these correlations were not statistically significant (Fig. S5A). After characterizing the individual effects of all single species, we assayed each pair of affecting species against the focal species. Overall, negative effects were significantly more prevalent in joint pair effects (73%) than in effects of individual species (59.4%) (p=3e-8, Fisher’s exact test) (Fig. 2B,C). Moreover, joint pair effects were quantitatively more negative than effects of individual species for most focal species on their own. The median effect on each focal was more negative by 0.49 on average, though the difference was not significant in most cases, and some focals showed a small increase in median effect (Fig. 2D). Despite this, the minimum and maximum effects for each focal species remained similar. As with the single affecting species, pairs’ joint effects did not correlate well with species traits, with similarity between the two affecting species, or with their similarity to the focal species (Fig. S5B). These results indicate that it may be challenging to connect the effects of single and pairs of species on a focal strain to a specific trait of the involved strains.

### Joint effects are not additive, but dominated by the stronger single species interaction

Next, we examined how the effects of individual species relate to their joint effect. In particular, we were interested in finding a model that describes the effects of pairs, based on the data from single species effects. Based on previous studies’ success in predicting community structure from pairwise interactions (Friedman, Higgins, and Gore 2017; Meroz et al. 2021; Guo and Boedicker 2016; Os et al. 2018), we posited that predicting how effects combine based solely on the effects of the single species should also be feasible. To do so, we considered three models: an additive effect model, a mean effect model, and a strongest effect model.

The additive effect model proposes that the effects of each species on the focal are the same whether they act individually or jointly. Therefore, the combined effect will be equal to the sum of the effects of each species on their own. This is equivalent to additivity of effects between antibiotics, which is common in the drug combinations field (Bollenbach 2015). The mean effect model assumes that the per capita effect of each species will remain the same, but that each species’ population size will be decreased and will be proportional to their initial fraction in the community. In contrast, the strongest effect model posits that the species with the strongest effect dominates the effect of other affecting species, leaving the joint effect the maximum single effect, and not the sum or mean of single species effects.

We found that the model that best agrees with the measured effects is the strongest effect model (Fig. 3). The strongest effect model was the most accurate overall (nRMSE=0.65, 0.47, 0.28 for the additive, mean, and strongest models, correspondingly). The strongest model was considerably more accurate when both species affected the focal negatively (nRMSE=0.66, 0.26, 0.18), slightly more accurate when both species affected the focal positively (nRMSE=0.63, 0.63, 0.53), and slightly less accurate when one effect was negative and the other positive (nRMSE=0.28, 0.62, 0.34) (Fig. S7A). Though supplementing the mean model with additional species information (i.e. carrying capacity) did improve the model accuracy, it was still less accurate than the strongest effect model (Fig. S6). The distribution of errors further supported the strongest effect model (Fig. 3B, Fig. S7B): When both single species effects were negative, the mean model was prone to underestimating the combined effect, due to the reduction of the stronger effect by taking into account the weaker effect; while contrastingly, the additive model overestimated effects due to the addition of the weaker effect to the stronger effect, which was more accurate on its own. We saw the opposite trend when both single species effects were positive, and no particular trend when there was one positive and negative effect. As with the effects themselves, model accuracy was not strongly correlated with any specific species trait (Fig. S8).

**Figure 3:**
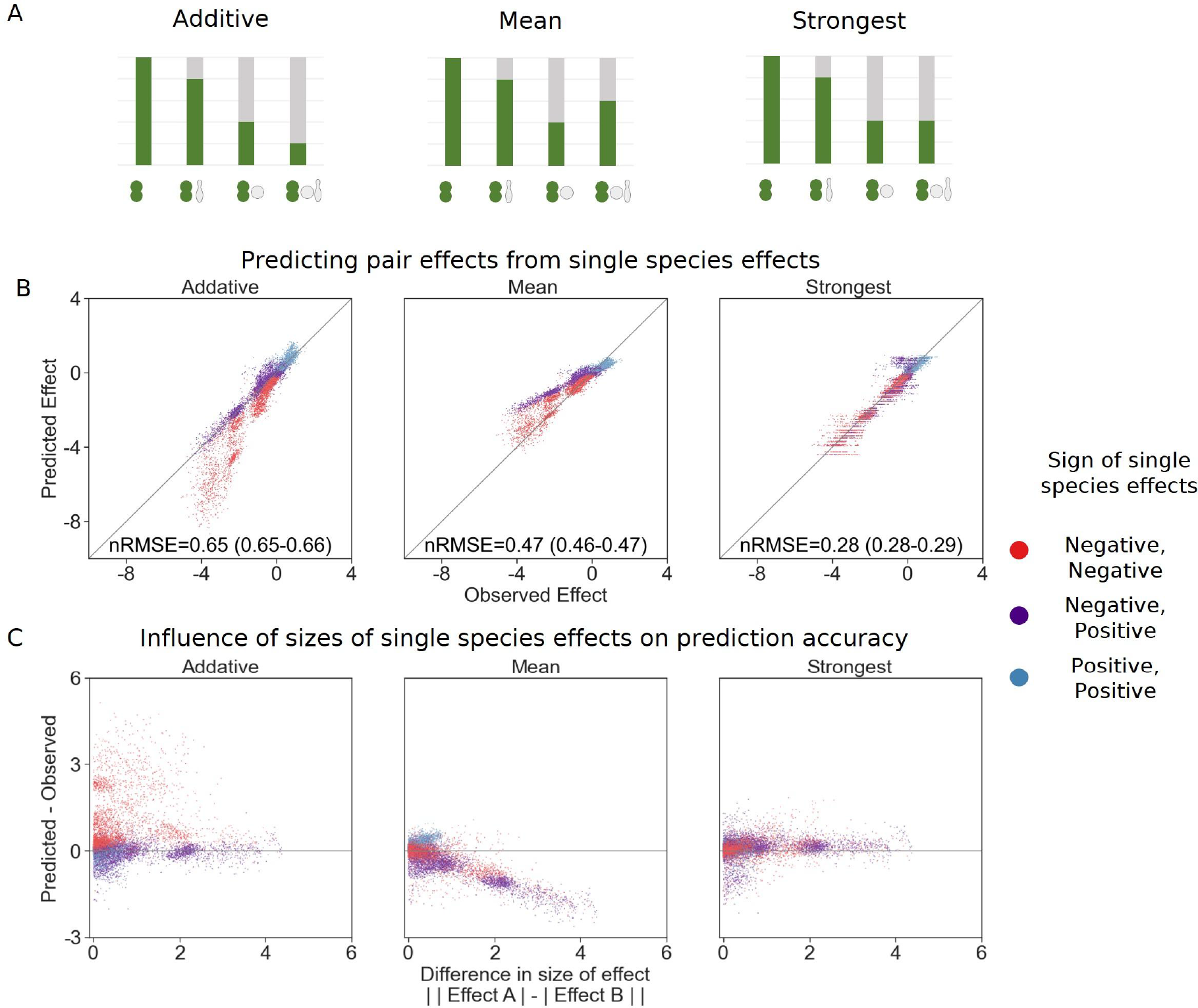
Strongest single species effect offers the most accurate model for the combined effect of two species. A) Graphical representation for each model. The additive model assumes that the effects of each species will accumulate, indicating they are acting independently, and are unaffected by one another. The mean model assumes the combined effect will be an average of the two single species effects. The final model, strongest effect, assumes that whichever species had a stronger effect on its own will determine the joint effect when paired with an additional species (see Materials and Methods for equations). B) Comparison of predicted effects and the experimental data, with their respective root mean squared error normalized to the interquartile range of the observed data (nRMSE). nRMSE values are calculated from 1000 bootstrapped datasets, and represent the median and interquartile range in parentheses (see Materials and Methods). Each dot represents the joint effect of a pair of affecting species on a focal species. Colors indicate the signs of the measured effects of the individual affecting species. C) The accuracy of each model as a function of the difference between the sizes of effect of each individual species within the pair.

Support for the strongest model is also evident in how the difference in size of effect influences the model accuracy (Fig. 3C). When effects are close to equal, the mean model is fairly accurate while the additive model does particularly poorly, as these effects would be calculated as twice the strongest effect. Contrastly, when one effect is much stronger than the other, the additive model is accurate since the addition of the weak effect is negligible, whereas the mean model underestimates the joint effects by taking into account the weaker effect.

### The strongest effect model is also the most accurate for larger communities

With this information in hand, we were interested to see if the same rules held up for larger communities. To this end, we screened trios of a subset (i.e. 30) of the affecting species against a single focal species (*E. coli*), and found similar trends to all those seen for pairs of affecting species. Similar to what was observed in the move from single species to pairs, there were more negative effects in trios than in the pairs (Fig. 4C). Additionally, as with joint pairs’ effects, the strongest effect model was more accurate than the additive and mean models. Similar distributions of error were seen as in the pairs’ effects, but further exaggerated with the more extreme under and over estimation of the combined trios’ effects by the mean and additive models respectively (Fig. 4A).

**Figure 4:**
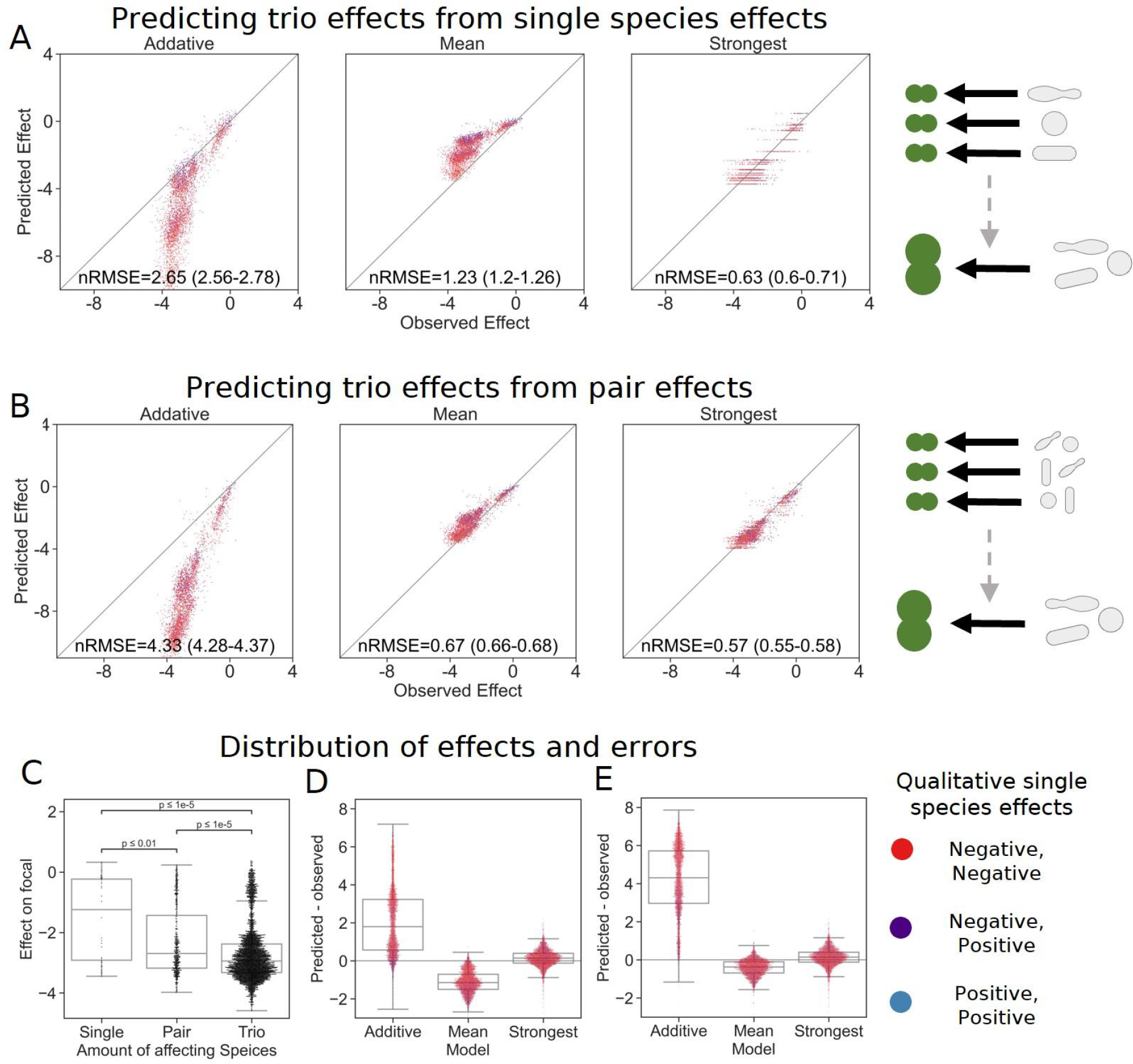
The strongest effect model is also the most accurate for trios. A,B) Correlation between three different models for how A) single species effects and B) pairwise species effects combine into trio effects, and the experimental data. Root squared mean error normalized to the interquartile range. nRMSE values are calculated from 1000 datasets, and represent the median and interquartile range in parentheses (see Materials and Methods). C) Distribution of the effects of single, pairs and trios of affecting species on E. coli. All Mann-Whitney-Wilcoxon two-sided tests were significant, p values are shown on plot. Dots show individual effects, solid lines represent the median, boxes represent the interquartile range, and whiskers are expanded to include values no further than 1.5X interquartile range. D,E) Distribution of errors for each model based on D) single species data and E) pairs data. Dots show individual effects, solid lines represent the median, boxes represent the interquartile range, and whiskers are expanded to include values no further than 1.5X interquartile range.

We further explored this model by basing trios’ data not on the additive, mean, or strongest values of the effects of individual species, but on those of the joint effects of the three pairs comprising each trio (the effects of single species and pairs were measured independently again in this experiment, see Materials and Methods, Fig. S1). The move to pairs-based predictions improved the accuracy for both the mean and strongest model, while further pushing the additive predictions away from the observed effects (Fig. 4B). These data suggest that even in the presence of additional species, the strongest single species effect still dominates the combined effect of a community.

## Discussion

By measuring over 14,000 simplified microbial communities, we quantified the effects of single species, pairs, and trios on multiple focal species. We observed that effects do not combine in an additive manner, but rather are dominated by the strongest effect of a single species. Despite most effects being negative, this is in stark contrast to models often used in antibiotic compound combinations where additivity is often the default model (Bollenbach 2015). The additive model performed well for mixed effects (i.e. one negative and one positive), but it seems unlikely this is due to effects remaining independent (as assumed in antibiotic combinations), as this model performed poorly when both species had effects of the same sign.

We phenotypically and genetically profiled all species, but did not find strong correlations between the measured traits, or similarity in traits, to the effect on the focal species. Though positive effects were common, making up about one third of the single species effects, they became less common as the number of community members increased, making up only one quarter of the effects of species pairs. Furthermore, we found similar trends in the larger communities of 4 species (three affecting species and one focal), both that effects combined in a non additive manner, being dominated by the strongest single species effect, and that negative effects became more common in larger communities which is consistent with previous studies (Cook et al. 2006; van Elsas et al. 2012; Jones et al. 2021; Piccardi, Vessman, and Mitri 2019; Gould et al. 2018; Palmer and Foster 2022).

It is difficult to identify a single biological mechanism that could explain why the strongest effect model agreed best with our experimental data. Intuitively, we assumed this could be explained by resource competition (i.e. an affecting species which consumes resources quickly would negatively affect the focal species, as well as the other affecting species). However, this explanation is not consistent with the fact that the affecting species’ growth rate did not correlate well with their effect on some focal species, and would not explain positive effects which act in a similar manner. Secondly, we thought effects could be saturating (either biologically, or with regard to the detection limits in this experimental setup), but this would not explain why the model works for weaker effects. It’s worth noting that a modified strongest effect model, which assumes mixed effects would always be negative (regardless of which is stronger), was more accurate for some focal species (Fig. S9), but this would also be poorly explained by saturating effects. A hierarchical ranking, where each species affects all the species ranked below it could lead to the strongest affecting species affecting both the focal and the other affecting species, thus dominating their joint effects, but this does not coincide with the fact that we observed almost no single species or pairs with a strong effect across all focals (Fig. S4). As we did not measure the presence of all species in each community (only the focal), we cannot disentangle interaction modification (changes in per capita effect of specific species), from interaction chains (affecting the amount of an affecting species, and as such its effect on the focal), and further work is needed in order to pinpoint the exact mechanism(s) leading to the dominance of the strongest effect in our system.

Understanding how microbial communities assemble, and how large numbers of species interact is of both utmost importance and difficulty. Harnessing such information would open up a plethora of currently underutilized applications in food, medical, and agricultural industries. Specifically, understanding how the effects of multiple species on a single species combine is important for introduction of new species into a given environment. Our results suggest that when we want to affect a single focal species in a given environment (e.g. for biocontrol of a pathogen), introducing the species with the strongest effect on the focal would be sufficient to obtain the desired effect, as synergies were rare in our dataset. In cases where there are multiple strains of interest (e.g. probiotics), introducing multiple species may be beneficial since different affecting species typically have strong effects on different focals. Introducing combinations of species may allow for a more robust function, as the chances that one member of the community will have a strong effect on a resident species of interest is more likely.

Further work is needed in order to deepen our understanding of how multiple species affect each other and to see to what extent our findings continue to hold up in more diverse communities, other taxonomic groups, and more complex environments. Nonetheless, our results suggest that community effects can be predicted from the strongest effect of a single species, greatly reducing the amount of information required to obtain accurate estimations, which can improve our ability to use a bottom-up approach for biotechnological applications, as well as answering fundamental ecology questions.

## Materials and Methods

### Strain isolation from soil samples

Soil (50 ml of soil, taken from a depth of ~30 cm) and leaf samples (multiple leaves from a single plant combined into a sterile 50 ml tube) were collected from various locations in the Faculty of Agriculture in Rehovot, Israel on multiple dates (See Data S1 file for more information). Each sample was diluted in phosphate-buffered saline (PBS) directly after collection (1 g of soil or one leaf in 10 ml of PBS) and vortexed for 5 minutes. 100 μl of multiple dilutions of this mixture (10^−2^-10^−5^) were seeded on different solid media [NB (0.5% [w/v] peptone, 0.3% [w/v] yeast extract, 1.5% [w/v] agar); 1% NB (0.005% [w/v] peptone, 0.003%[w/v] yeast extract, 1.5% agar); M9 minimal media (0.1 mM CaCl_2_, 2mM MgSO_4_, 1x [Enco-teknova] trace metals, 1% [w/v] glucose, 1x (Sigma) M9 Salts), additional plates were made with the same media containing various antibiotics (antibiotics and respective concentrations are listed in table S2) Plates were incubated at 30°C, and colonies were restreaked on NB without antibiotics until single isolates were stably obtained. Strains were selected on the basis of multiple criteria: growth of transferred colony in NB liquid medium (30°C), frozen glycerol stock revival in NB (OD_600_ > 0.1) (30°C), and subsequent growth on M9 minimal media + 1% (w/v) glucose (OD_600_ > 0.1) (30°C). Isolates were kept in single tubes as well as 96 well plates in 50% NB + 50% glycerol (glycerol stock were 60% and 80% for tubes and plates respectively for 30% and 40% final glycerol concentrations).

### Strain identification and phylogenetic distance calculation

Each bacterial isolate was classified phylogenetically with its 16S rRNA gene sequence. The full 16S gene sequences (~1500 base pairs) were obtained via Sanger sequencing, and classified with a combination of RDP Classifier(Cole et al. 2014) and BLAST(Altschul et al. 1990) (List of strains in table S1, full phylogenetic data in Data S1 file). Phylogenetic distance was calculated in Geneious Prime softwareֱֱ (version 2022.2.1, Biomatters Ltd). Sequences were aligned using MUSCLE alignment. Phylogenetic tree was built using the UPGMA method with no outgroup and a HKY genetic distance model. The pairwise phylogenetic distances between strains were calculated directly from the patristic distances of the phylogenetic tree.

### Phenotypic profiling and distance calculation

Bacterial strains were seeded from −80 stock directly into 1 ml LB medium (1% [w/v] tryptone, 1% [w/v] NaCl, 0.5% [w/v] yeast extract) in 96 well plate, and grown overnight at 30°C at 900 RPM (on a Titramax 100). Cells were washed 3 times by centrifugation as 3600 rfc, removal of supernatant, and resuspension in M9 minimal media with no carbon. Cultures were then normalized to 0.01 OD_600_. 20 μl of the normalized cultures were added to 180 μl of M9 minimal media either containing 1% [w/v] of one of 20 carbon sources (table S2) or M9 minimal media with 1% [w/v] glucose and one of 11 antibiotics (antibiotics and respective concentrations are listed in table S2). Plates were grown at 30°C for 48 hours without shaking. Cultures were homogenized by shaking (on a Titramax 100) for 90 seconds before measuring OD_600_. Additionally, species were also grown in M9 minimal media (with the addition of 0.05% [w/v] BSA and 1% [w/v] glucose) with shaking (continuous double orbital shaking, 282 cpm) at 30°C, with OD measurements every 15 minutes, for 48 hours, to obtain growth kinetics (in Epoch and Synergy H1 microplate readers). Growth for carbon source experiments was normalized to the carbon source with the highest OD_600_ and antibiotic experiments were normalized to growth on M9 minimal media with glucose and no antibiotics. Euclidean distances of normalized values were measured for each species on carbon sources and antibiotics separately, and used to construct distance matrices.

### Droplet preparation and culturing

Bacterial strains were seeded from −80 stock directly into 1 ml LB medium (1% [w/v] tryptone, 1% [w/v] NaCl, 0.5% [w/v] yeast extract) in 96 well plate, and grown overnight at 30°C at 900 RPM (on a Titramax 100). Cells were washed 3 times by centrifugation as 3600 rfc, removal of supernatant, and resuspension in M9 minimal media (with the addition of 0.05% [w/v] BSA and 1% [w/v] glucose). Affecting species cultures were then normalized to 0.04 OD_600_ and focal species cultures were to 0.02 OD_600_. Affecting and focal species cultures were combined at a ratio of 1:1 so that droplets contained a final concentration of 0.02 OD_600_ affecting species and 0.01 OD_600_ focal species. (With this setup, in a well with two distinct droplets, there is a starting concentration of 0.01 OD_600_ for each species.)

Droplets were produced on a Bio-Rad QX200 Droplet Generator as described by Kulesa *et al*.(Kulesa et al. 2018) Briefly, 20 μl input of combined cultures were emulsified into ~20,000 1 nl droplets in fluorocarbon oil (3M Novec 7500) stabilized with 2% (w/w) fluorosurfactant (RAN Biotech 008 FluoroSurfactant). 2.5 mM of fluorescent dyes (Thermo Fisher AlexaFluor™: 555 (A33080), 594 (A33082), 647 (A33084)) were added to culture for droplet imaging (See Kulesa et al.(Kulesa et al. 2018)). For each kChip loading, about 5000 droplets for each input (~60 affecting species + focal species, 2 focal species monocultures, 2 blank cultures) were generated, for a total of ~320,000 droplets. Droplets were generated together for 2 kChips (technical replicates), and then droplets were pooled separately for each chip. kChips were incubated at 30°C for 72 hours. Cultures were imaged at 24 hour intervals throughout the experiment. Data for analysis was taken from after 24 hours, as monoculture growth of the focals saturated by this point (Fig. S10).

### Data filtering and normalization

As the kChip genreates droplet combinations stochastically, the amount of replicates for each community is different, ranging from 1 to 285, with a mean of 19. All communities with less than 3 replicates were not used in the analysis. Normalization was performed by subtracting the starting value for each individual well from the additional time points.

### Calculating effect size

To measure the effect of each affecting species on a given focal species, the log of the ratio of focal species yield in coculture (median of coculture replicates) to monoculture (median of monoculture replicates) was calculated:

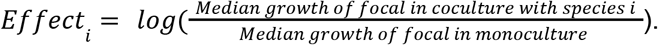

Coculture data was collected from wells with different starting concentrations in both the two-droplet experiments (i.e. one mixed droplet and one focal monoculture) and three-droplet experiments (i.e. two droplets of affecting species A and one droplet of affecting species B and vice versa, or one of each in addition to a blank droplet) (Fig. S1). Our data showed that the different initial fractions did not influence the effect on the focal species (Fig. S11). The standard error was calculated via bootstrapping, 100 calculations of the resampled median coculture divided by resampled median monoculture. Effects where the standard deviation was larger than the absolute value of the effect were classified as neutral.

### Calculating predictions for different models and their accuracy

The additive model assumes that the effects of each species will accumulate, and is the combined effect is the sum of effects, calculated as:

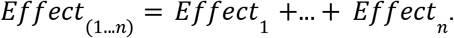

The mean model assumes the combined effect will be an average of the two single species effects, and is calculated as:

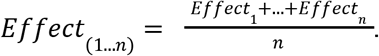

The OD-weighted mean model weighs the mean of effects by the affecting species’ maximum OD_600_ in the growth curves experient and is calculated as:

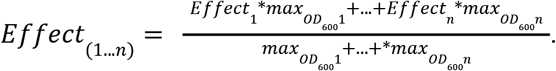

The strongest effect model assumes that whichever species had a stronger effect on its own will determine the joint effect when paired with an additional species. It is calculated as:

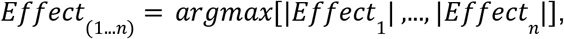

returning the effect with the largest absolute value (e.g. if two single species’ effects are −3 and +1, the model will predict that their joint effect is −3).

Root mean square error measuring the accuracy of each model was normalized to the interquartile range for each dataset. Normalized root mean square error was calculated via bootstrapping, 1000 calculations of the resampled wells from each chip were used to calculate the median and interquartile range of the normalized root mean square errors.

## Acknowledgements

We thank Jared Kehe and Anthony Ortiz for helping set up the kChip system in our lab, as well as Meilin Zhu, Megan Tse, Julie Chen and Paul Blainey for continued correspondence and technical support. Alfonso Pérez Escudero, and members of the Friedman lab for helpful discussions. Nadav Kashtan for constructive comments on the manuscript. Eddie Cytryn for generously providing bacterial strains used in this study. Lastly, Yael Sorokin for technical assistance. This research was supported by the United States – Israel Binational Science Foundation (grant no. 2017179).

## Author information

Authors Affiliations:

Amichai Baichman-Kass, Tingting Song, & Jonathan Friedman

Institute of Environmental Sciences, Hebrew University, Rehovot, Israel

Author Contributions:

A.B.K. and J.F. designed the study and wrote the manuscript, ABK performed the experiments, and analysis. T.S. isolated strains used in this study.

## Notes

### Competing Interest Statement

The authors have declared no competing interest.

### Summary of Updates

We have added a bootstrap analysis for nRMSEs presented in the paper. We have added accession numbers for strains isolated in this study to Data S1 file. Minor syntax revisions.

