## Supplementary Materials for "Interactions between culturable bacteria are highly non-additive"

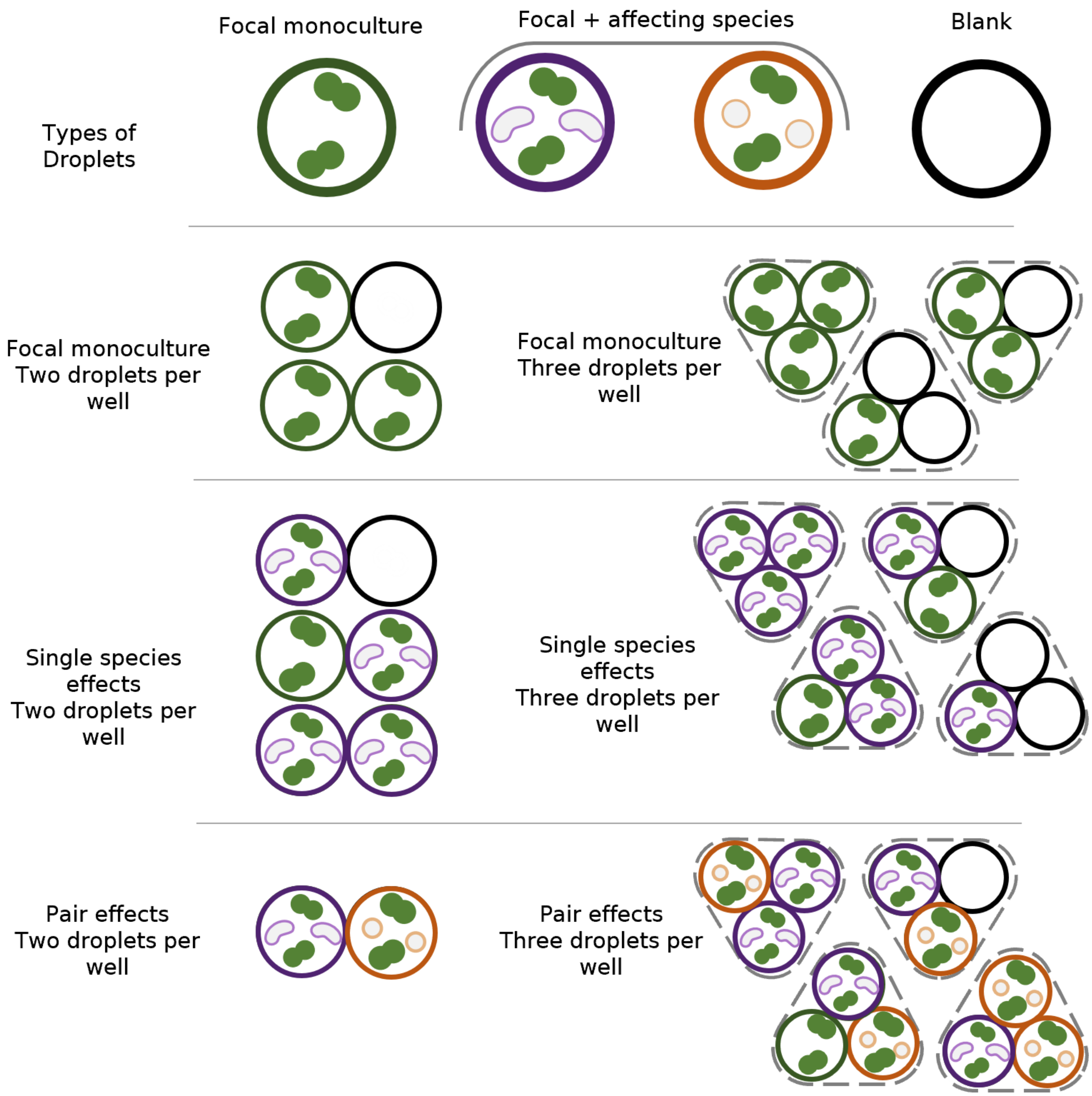


**Figure S1**: **Species in each droplet and well in kChip experiments**

*Droplets contained one of three different types of samples: 1) focal monoculture, containing only the focal species, 2) coculture of focal species and a single affecting species at equal optical densities, 3) Blank samples, with media but no bacterial cells. Based on which two or three droplets (depending on the experimental setup) each well contained either a focal monoculture, one, two or three, affecting species. Similar communities started at different initial densities based on how many monoculture or blank droplets were in the well, but as shown in Fig. S11, initial densities did not influence the effect on the focal.*

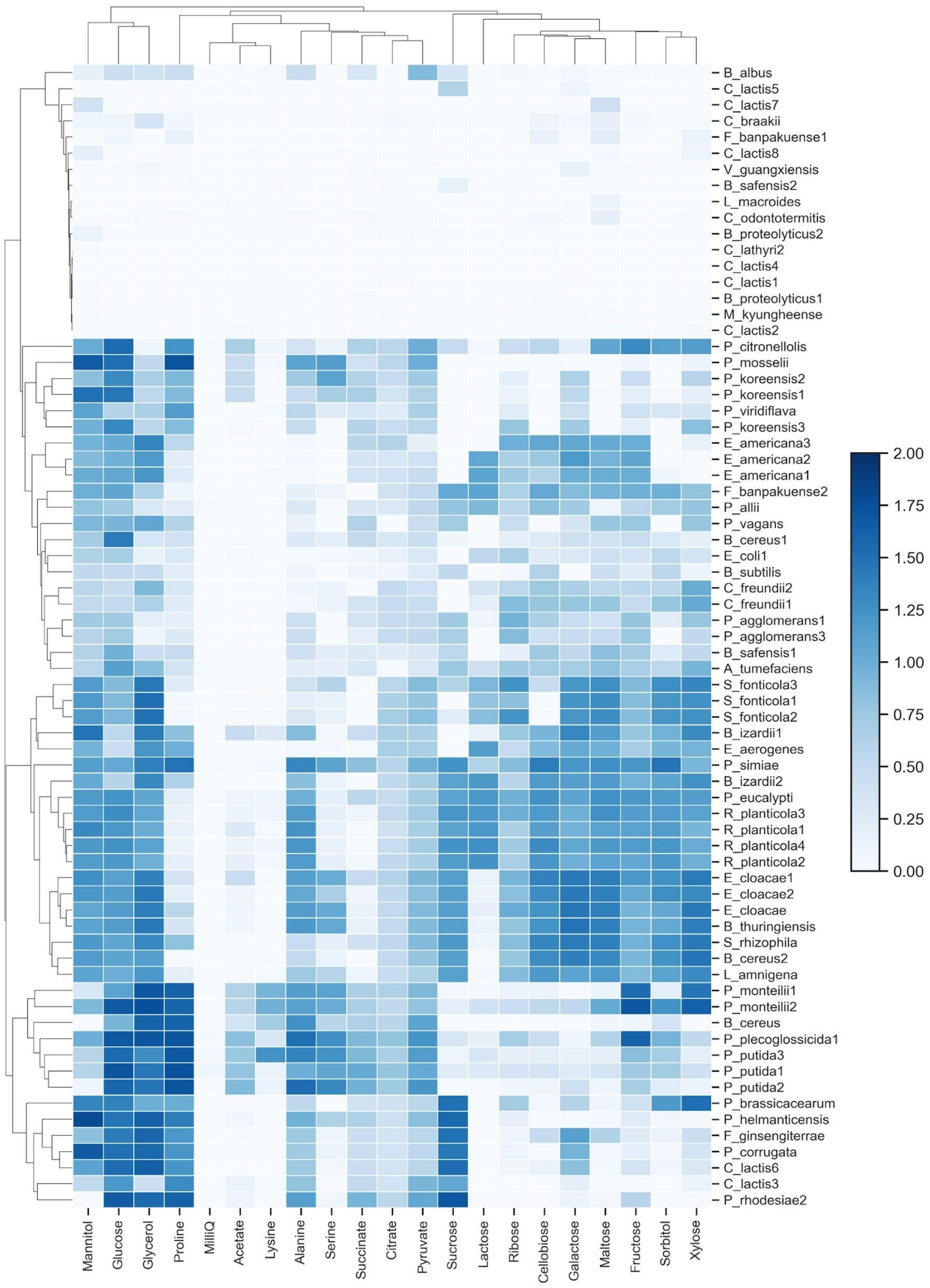


**Figure S2**: **Carbon source utilization profiles for bacterial strains**

*Each strain's yield on 20 different carbon sources, assayed after 48 hours. Growth values are calculated as mean $OD_{600}$ measurement from 3 replicates, and were background-subtracted (media with no bacteria).*

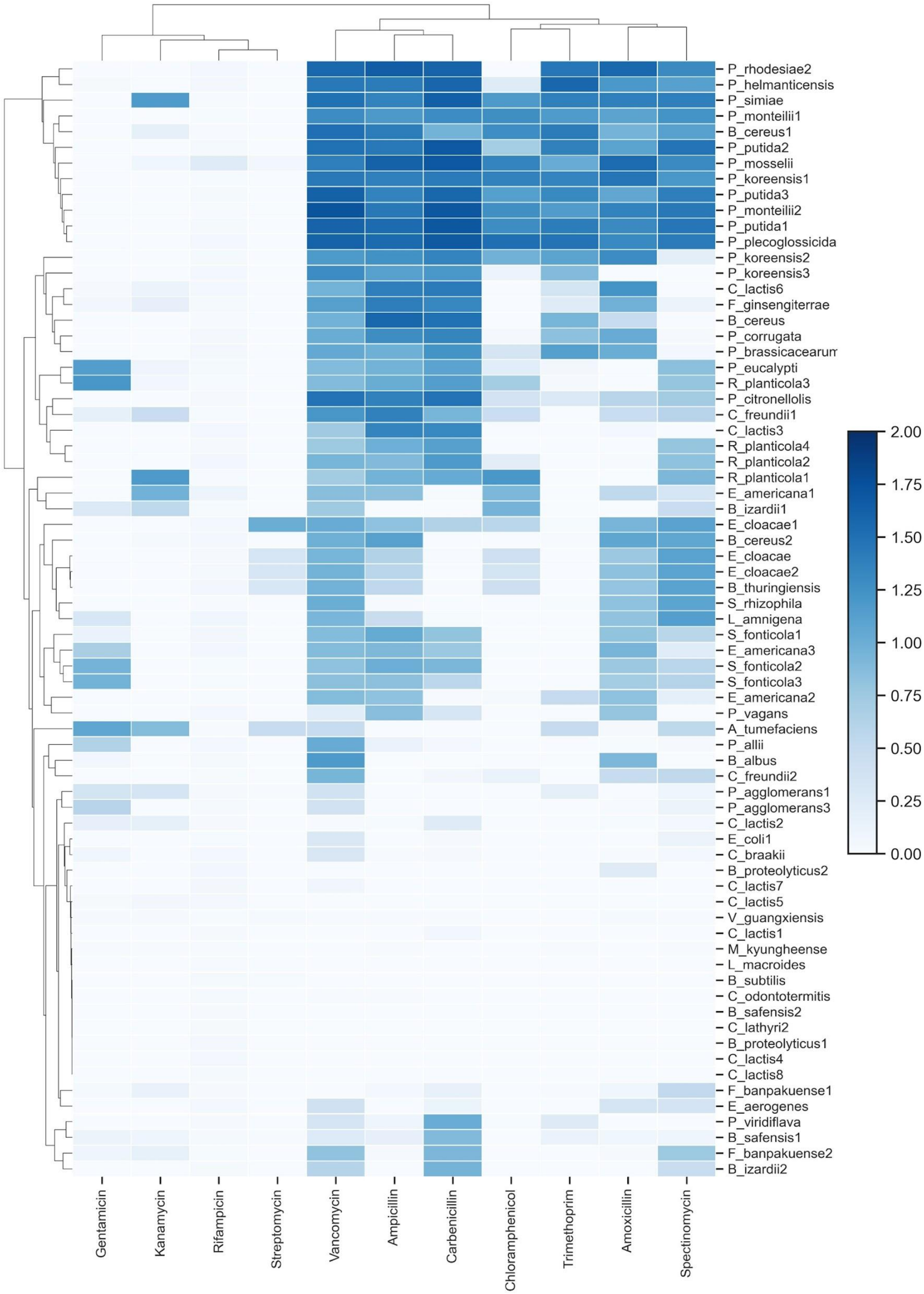


**Figure S3**: **Antibiotic resistance profiles for bacterial strains**

*Each strain's ability to grow on 11 different antibiotics, after 48 hours. Growth values are calculated as mean $OD_{600}$ measurement from 3 replicates, and were background-subtracted (media with no bacteria).*

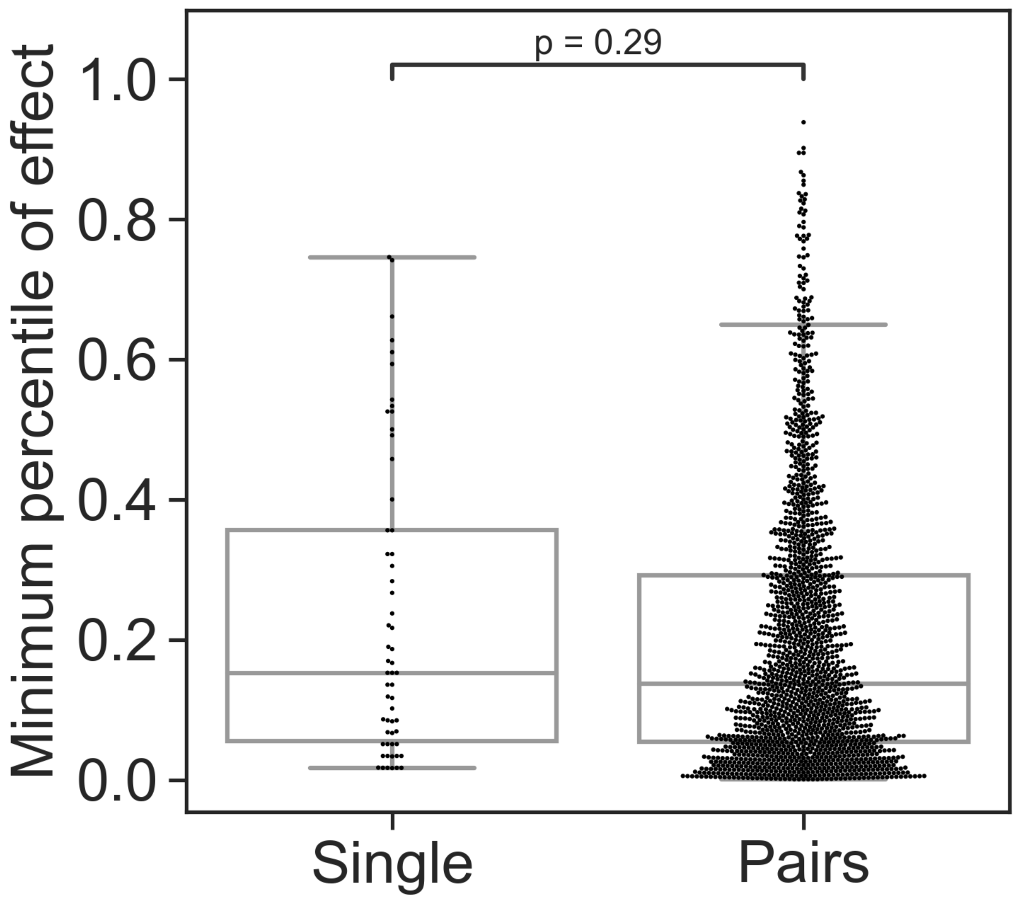


**Figure S4**: **Minimal effect of single strains and pairs across multiple focals**
*Distributions of the weakest effect of each individual species and pairs measured as the percentile within the distributions of effects for a single focal species. For each species or pair, the minimal value from all focals was taken to generate the above distributions. Sign was not regarded in this calculation, only strength of the effect. Dots represent individual measurements, solid lines represent the median, boxes represent the interquartile range, and whiskers are expanded to include values no further than 1.5X interquartile range. Mann-Whitney-Wilcoxon test two-sided tests were performed,and p-value is shown on the graph.*

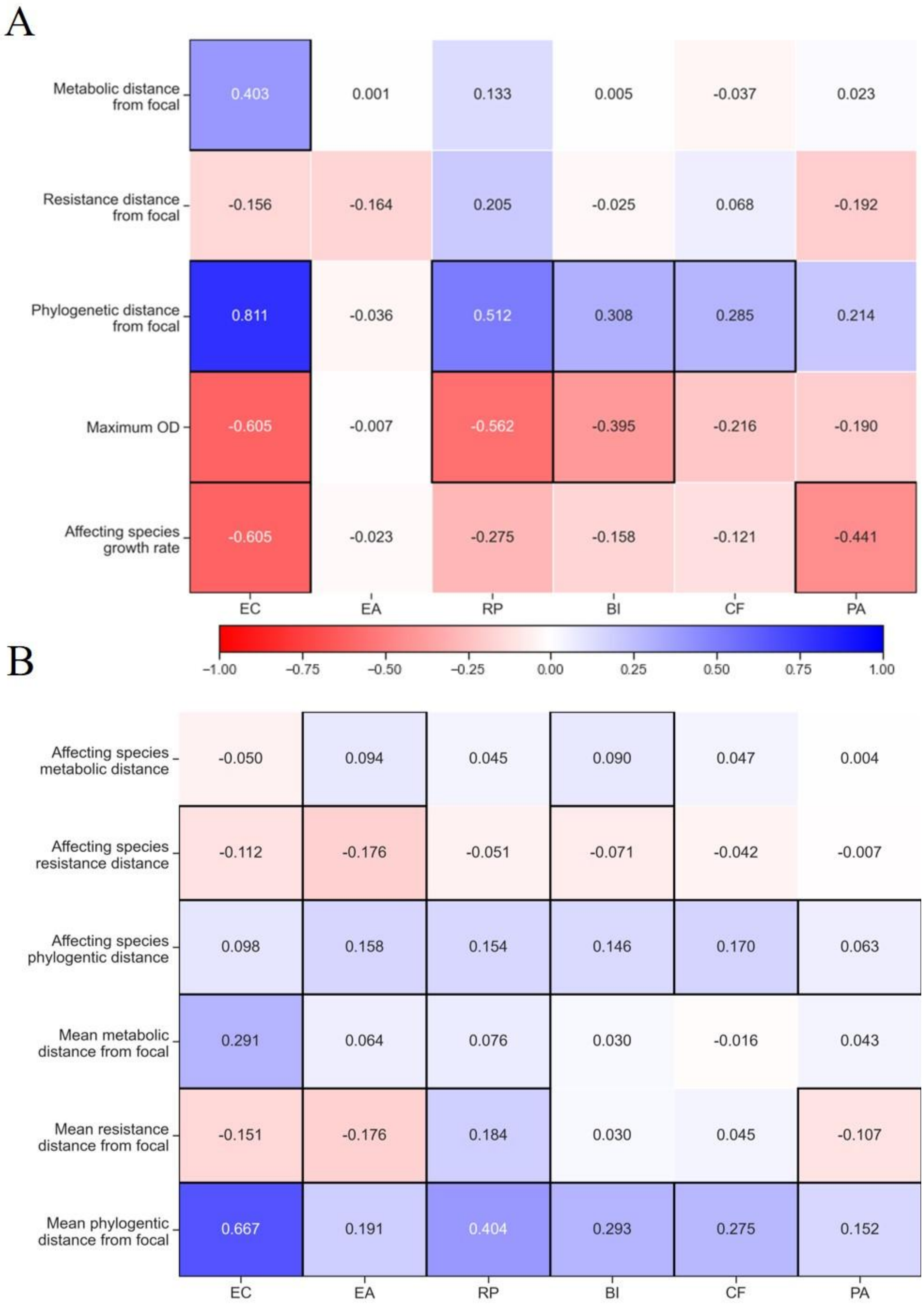


**Figure S5**: **Correlation between affecting species traits and effect on focal**

*Pearson correlation value for traits of A) single species, and B) pairs and effect on species separated for each focal species individually. Correlations with p-values below 0.05 are highlighted with a black frame. See Materials and Methods for calculations of phenotypic and phylogenetic distances.*

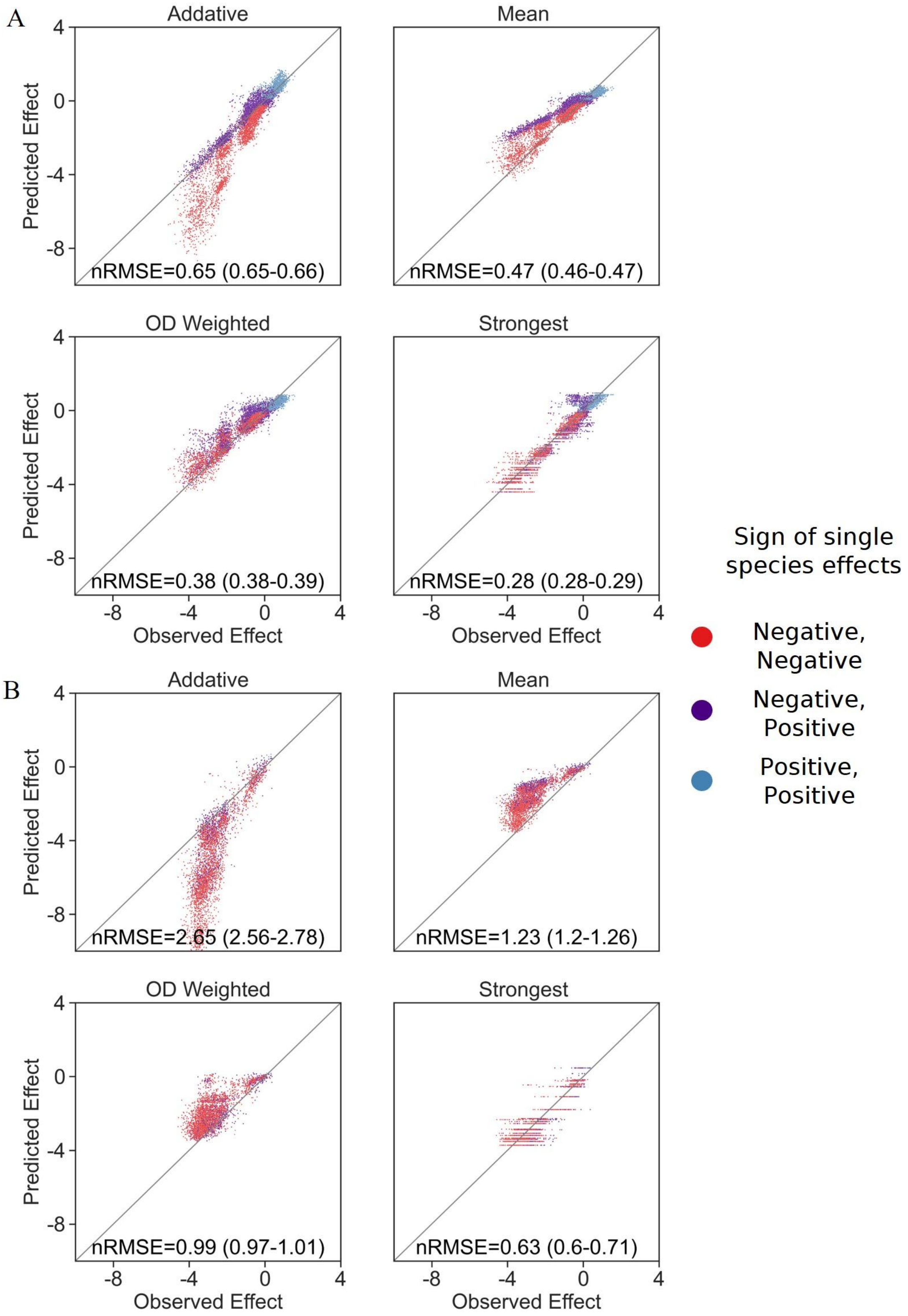


**Figure S6**: **OD weighted mean model**

*Correlation between four different models for how single species effects combine into A) pair effects and B)trio effects, and the experimental data, with their respective normalized root mean squared error. nRMSE values are calculated from 1000 bootstrapped datasets, and represent the median and interquartile range in parentheses (see Materials and Methods). Similar to Fig. 3B with the addition of the OD weighted mean.*

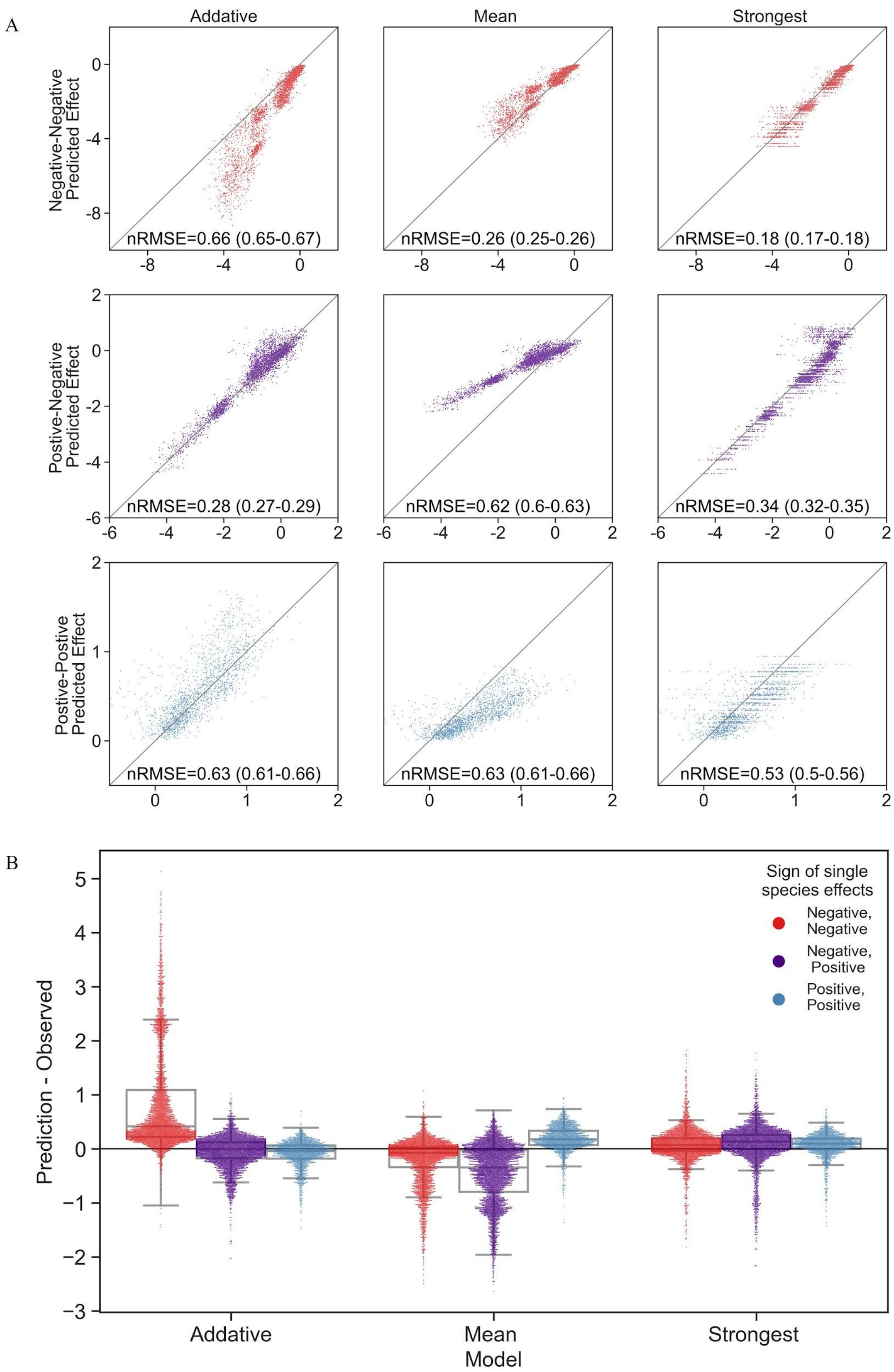


**Figure S7**: **Distribution of errors for each model predicting pair effects from single species**

A) *Correlation between three models for how single species effects combine, and the experimental data, with their respective normalized root squared mean error. nRMSE values are calculated from 1000 bootstrapped datasets, and represent the median and interquartile range in parentheses (see Materials and Methods). Data is divided by interaction signs of the individual species.* B) *Distribution of the prediction accuracy for each model. Dots represent individual measurements, solid lines represent the median, boxes represent the interquartile range, and whiskers are expanded to include values no further than 1.5X interquartile range. The frequencies of these interaction types in the dataset are Negative-Negative 39%, Positive-Positive 18% and Negative-Positive 43%.*

.

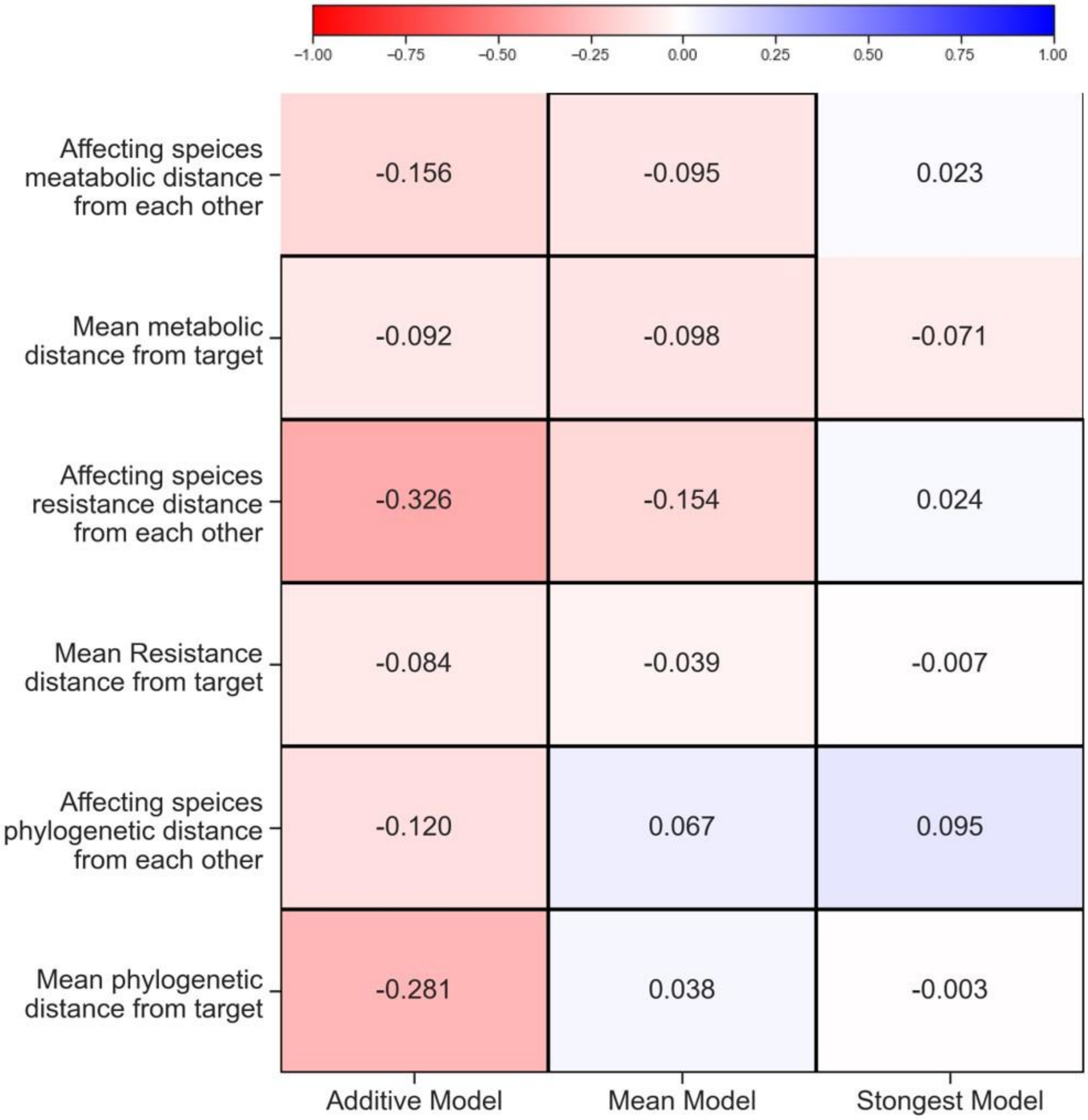


**Figure S8**: **Traits effect on model error**

*Pearson correlation value for each trait and the deviation of the model. Correlations with p-values below 0.05 are highlighted with a black frame. See see Materials and Methods for calculations of phenotypic and phylogenetic distances*

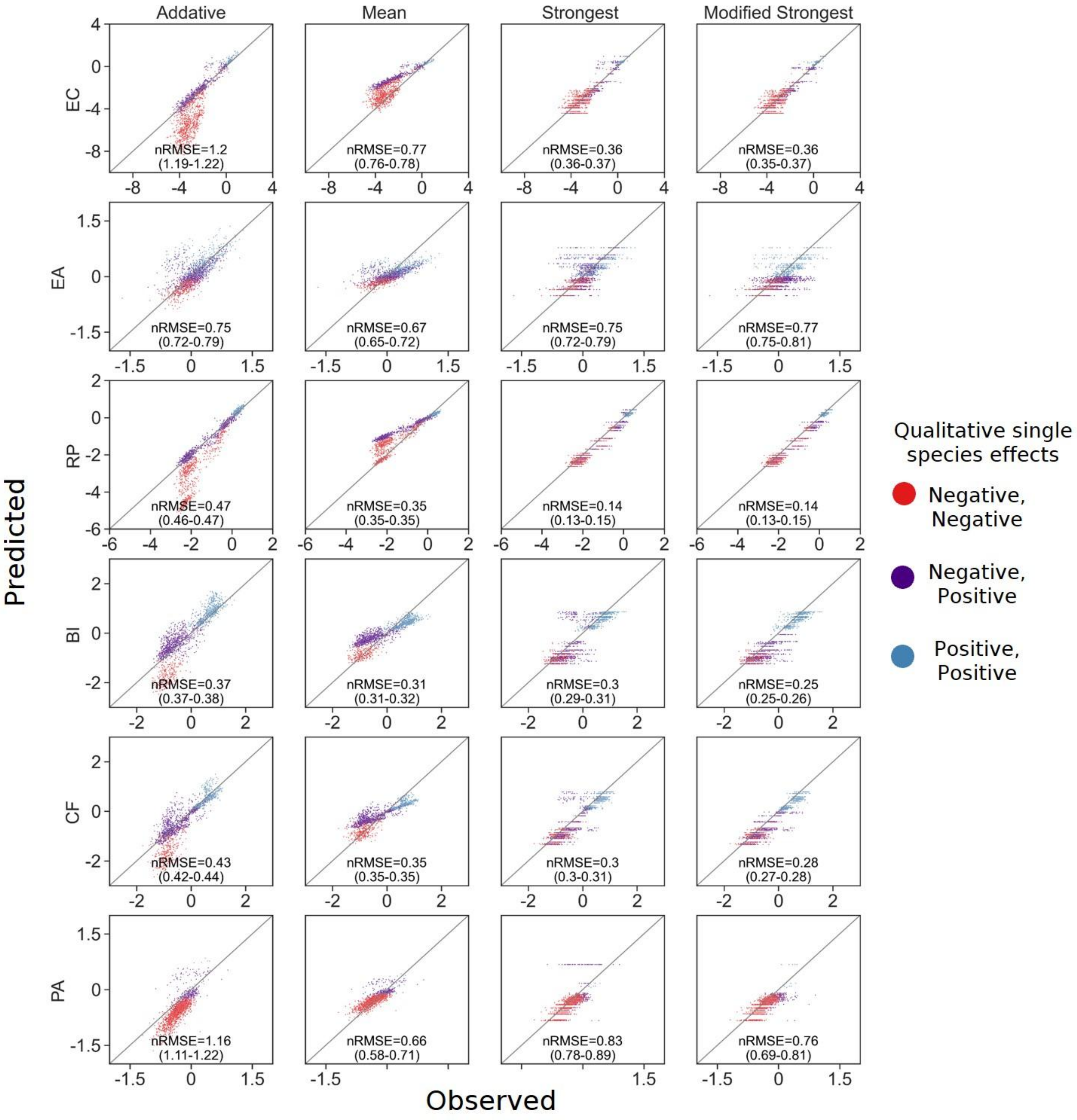


**Figure S9**: **Model comparisons divided by focal species**

*Correlation between four different models for how single species effects combine, and the experimental data, with their respective normalized root squared mean error.nRMSE values are calculated from 1000 bootstrapped datasets, and represent the median and interquartile range in parentheses (see Materials and Methods). Data is divided for each focal species individually. In addition to the three models explained in detail in the main text, a modified strongest effect model, which assumes mixed effects would always be negative (regardless of which is stronger), was added to this analysis as well.*

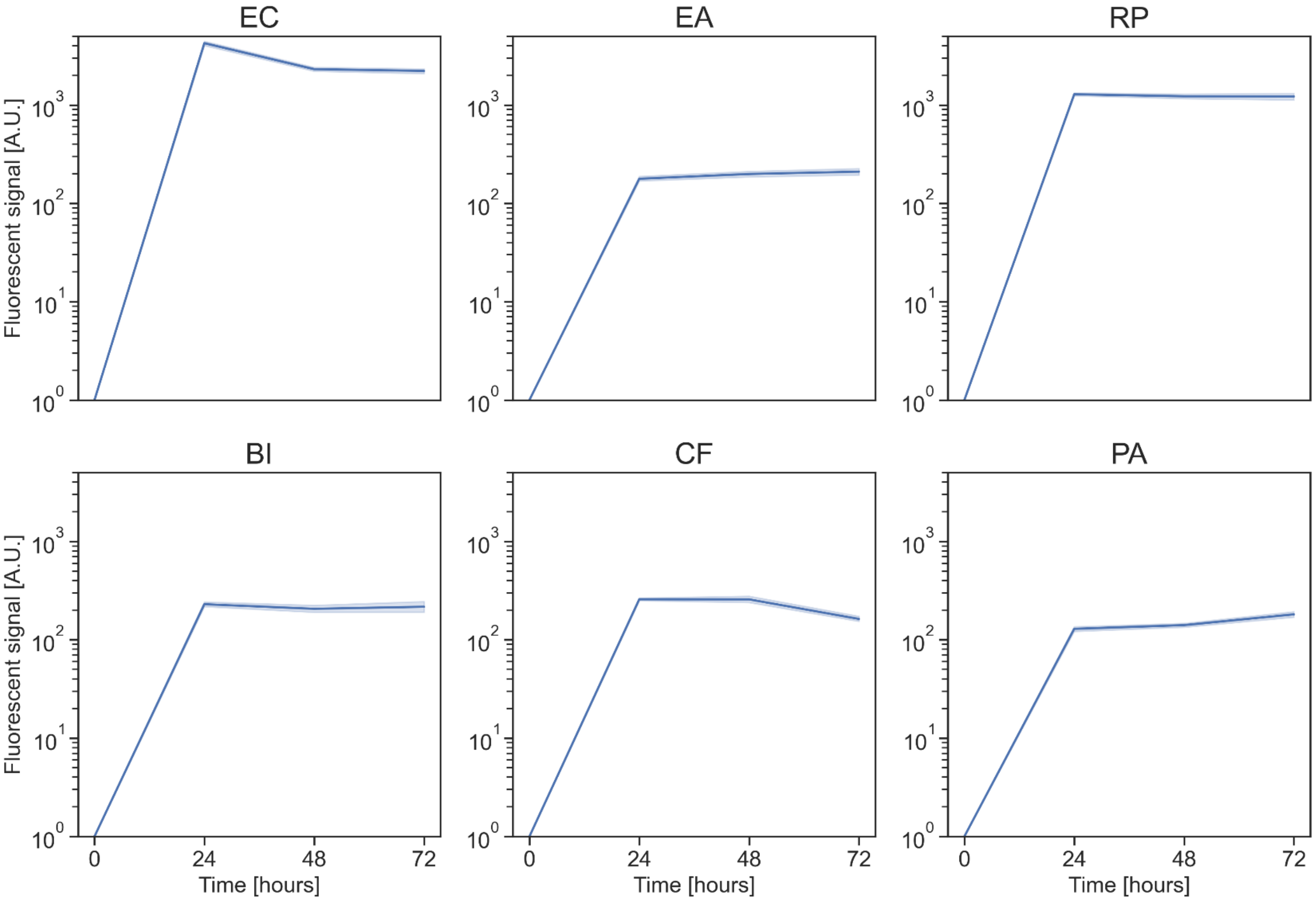


**Figure S10**: **Growth dynamics of focal species in monoculture**

*Growth in kChip of each focal strain in monoculture over 72 hours. Growth is measured by fluorescent signal and each well is normalized to the value at the beginning of the experiment (all signals at time zero are equal to one). The solid line represents the mean, and the shaded area the 95% confidence interval.*

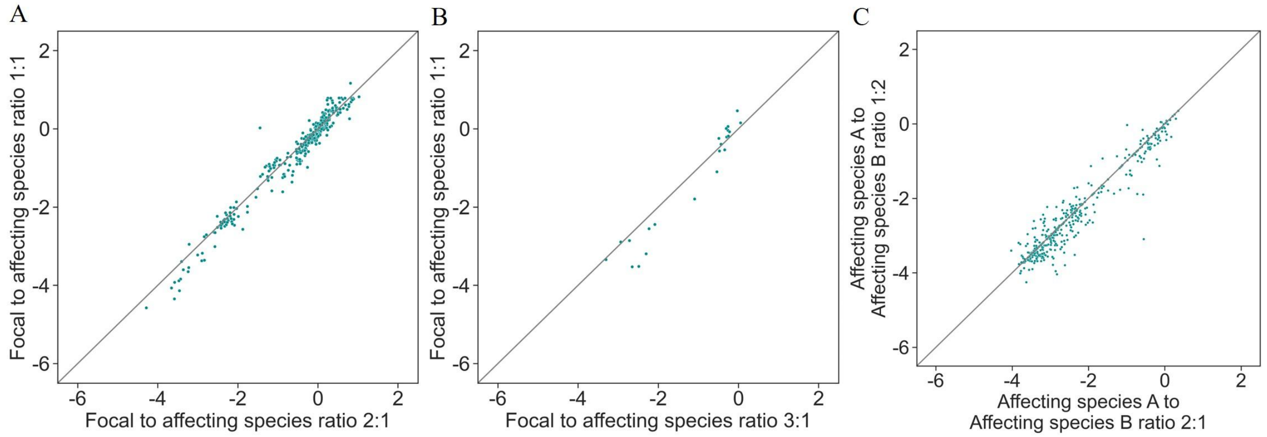


**Figure S11**: **Effects on focal species are independent of initial species' density**

*Correlation between different ratios of initial optical density in each well. A) Different ratios of focal to affecting species based on whether there were two droplets containing the affecting species or one and one focal monoculture droplet-* $R^2$*=0.96. B) Different ratios of focal to affecting species based on whether there were three droplets containing the affecting species or one and two focal monoculture droplets-* $R^2$*=0.94. C) Different ratios of affecting species based on whether there were two droplets containing affecting species A and one droplet containing affecting species B or vice versa-* $R^2$*=0.92. All p-values were below 1e-10.*

**S1 Table 1: Strains used in this study**

Rows highlighted in green are represent the focal species, full phylogenetic data included in Data S1 file

| Nearest Species | Source | In Trios Subset | | Nearest Species | Source | In Trios Subset |
|---|---|---|---|---|---|---|
| Microbacterium kyungheense | This Study | Yes | | Ewingella americana | [20] | Yes |
| Chryseobacterium lactis | This Study | | | Ewingella americana | [20] | |
| Chryseobacterium lactis | This Study | Yes | | Ewingella americana | [20] | |
| Chryseobacterium lactis | This Study | Yes |  | Lelliottia amnigena | [20] |  |
| Chryseobacterium lactis | This Study | Yes | | Pantoea agglomerans | [20] | |
| Chryseobacterium lactis | This Study | Yes | | Pantoea agglomerans | [20] | |
| Chryseobacterium lactis | This Study | Yes | | Pantoea agglomerans | [20] | |
| Chryseobacterium lactis | This Study | Yes | | Pantoea allii | [20] | |
| Chryseobacterium lactis | This Study | | | Pantoea eucalypti | [20] | |
| Chryseobacterium lathyri | This Study | Yes | | Pantoea vagans | [20] | |
| Chryseobacterium lathyri | This Study | | | Pseudomonas brassicacearum | This Study | |
| Flavobacterium banpakuense | This Study | Yes | | Pseudomonas citronellolis | This Study | |
| Flavobacterium banpakuense | This Study | Yes | | Pseudomonas corrugata | This Study | Yes |
| Flavobacterium ginsengiterrae | This Study | Yes | | Pseudomonas helmanticensis | [20] | |
| Bacillus albus | This Study | Yes | | Pseudomonas koreensis | [20] | Yes |
| Bacillus cereus | This Study | Yes | | Pseudomonas koreensis | This Study | Yes |
| Bacillus cereus | Cytryn Lab | | | Pseudomonas koreensis | This Study | |
| Bacillus cereus | Cytryn Lab | | | Pseudomonas monteilii | This Study | Yes |
| Bacillus proteolyticus | Cytryn Lab | | | Pseudomonas monteilii | This Study | Yes |
| Bacillus proteolyticus | Cytryn Lab | | | Pseudomonas mosselii | This Study | Yes |
| Bacillus safensis | This Study | | | Pseudomonas plecoglossicida | [20] | |
| Bacillus safensis | This Study | | | Pseudomonas putida | This Study | Yes |
| Bacillus subtilis | Cytryn Lab | | | Pseudomonas putida | This Study | Yes |
| Bacillus thuringiensis | Cytryn Lab | | | Pseudomonas putida | This Study | Yes |
| Lysinibacillus macroides | This Study | Yes | | Pseudomonas rhodesiae | [20] | |
| Agrobacterium tumefaciens | [20] | | | Pseudomonas rhodesiae | [20] | |
| Buttiauxella izardii | [20] | | | Pseudomonas simiae | This Study | Yes |
| Buttiauxella izardii | [20] | Yes | | Pseudomonas viridiflava | This Study | |
| Citrobacter braakii | [20] | | | Raoultella planticola | [20] | |
| Citrobacter freundii | [20] | | | Raoultella planticola | [20] | Yes |
| Citrobacter freundii | [20] | | | Raoultella planticola | [20] | |
| Comamonas odontotermitis | This Study | | | Raoultella planticola | [20] | |
| Enterobacter aerogenes | [20] | | | Serratia fonticola | [20] | |
| Enterobacter cloacae | Cytryn Lab | Yes | | Serratia fonticola | [20] | |
| Enterobacter cloacae | Cytryn Lab | | | Serratia fonticola | [20] | |
| Enterobacter cloacae | Cytryn Lab | | | Stenotrophomonas rhizophila | This Study | Yes |
| Escherichia coli | [20] | Yes | | Variovorax guangxiensis | This Study | |
| Escherichia coli | [20] | | | | | |

SI Table 2: Carbon sources and antibiotics used in species profiling experiment

| Carbon Source | Conc. | Antibiotic | Conc. |
|---|---|---|---|
| Glucose D+ | 1% w/v | Kanamycin | 50 µg/ml |
| D-Ribose | 1% w/v | Spectinomycin | 60 µg/ml |
| D-Xylose | 1% w/v | Amoxicillin | 50 µg/ml |
| D-Fructose | 1% w/v | Gentamicin | 20 µg/ml |
| D-Galactose | 1% w/v | Streptomycin | 100 µg/ml |
| Sucrose | 1% w/v | Trimethoprim | 50 µg/ml |
| D-Cellobiose | 1% w/v | Carbenicillin | 12.5 µg/ml |
| D-Lactose monohydrate | 1% w/v | Vancomycin | 50 µg/ml |
| D-Maltose monohydrate | 1% w/v | Rifampicin | 100 µg/ml |
| Sodium Citrate Dihydrate | 1% w/v | Chloramphenicol | 20 µg/ml |
| Disodium Succinate | 1% w/v | Ampicillin | 100 µg/ml |
| Sodium Acetate | 1% w/v | | |
| Sodium Pyruvate | 1% w/v | | |
| Glycerol | 1% v/v | | |
| D-Mannitol | 1% w/v | | |
| D-Sorbitol | 1% w/v | | |
| L-Serine | 1% w/v | | |
| L-Alanine | 1% w/v | | |
| L-Proline | 1% w/v | | |
| L-Lysine | 1% w/v | | |
